# Foundations of human problem solving

**DOI:** 10.1101/779322

**Authors:** Noah Zarr, Joshua W. Brown

## Abstract

Despite great strides in both machine learning and neuroscience, we do not know how the human brain solves problems in the general sense. We approach this question by drawing on the framework of engineering control theory. We demonstrate a computational neural model with only localist learning laws that is able to find solutions to arbitrary problems. Using a combination of computational neural modeling, human fMRI, and representational similarity analysis, we show here that the roles of a number of brain regions can be reinterpreted as interacting mechanisms of a control theoretic system. The results suggest a new set of functional perspectives on the orbitofrontal cortex, hippocampus, basal ganglia, anterior temporal lobe, lateral prefrontal cortex, and visual cortex, as well as a new path toward artificial general intelligence.

---

Great strides have been made recently toward solving hard problems with deep learning, including reinforcement learning (*1, 2*). While these are groundbreaking and show superior performance over humans in some domains, humans nevertheless exceed computers in the ability to find creative and efficient solutions to novel problems, especially with changing internal motivation values. Artificial general intelligence (AGI), especially the ability to learn autonomously to solve arbitrary problems, remains elusive (*3*).

Value-based decision-making and goal-directed behavior involve a number of interacting brain regions, but how these regions might work together computationally to generate goal directed actions remains unclear. This may be due in part a lack of mechanistic theoretical frameworks (*4, 5*). The orbitofrontal cortex (OFC) may represent both a cognitive map (*6*) and a flexible goal value representation (*7*), driving actions based on expected outcomes (*8*), though how these guide action selection is still unclear. The hippocampus is important for model-based planning (*9*) and prospection (*10*), and the striatum is important for action selection (*11*). Working memory for visual cues and task sets seems to depend on the visual cortex and lateral prefrontal regions, respectively (*12, 13*).

Neuroscience continues to reveal aspects of how the brain might learn to solve problems. Studies of cognitive control highlight how the brain, especially the prefrontal cortex, can apply and update rules to guide behavior (*14, 15*), inhibit behavior (*16*), and monitor performance (*17*) to detect and correct errors (*18*). Still, there is a crucial difference between rules and goals. Rules define a mapping from a stimulus to a response (*19*), but goals define a desired state of the individual and the world. When cognitive control is re-conceptualized as driving the individual to achieve a desired state, or set point, then cognitive control becomes a problem amenable to control theory. Control theory has successfully accounted for the neural control of movement (*20*) but has not been applied to cognition more broadly. In this framework, a preferred decision prospect will define a set point, to be achieved by control-theoretic negative feedback controllers (*21, 22*). Problem solving then requires 1) defining the goal state; 2) planning a sequence of state transitions to move the current state toward the goal; and 3) generating actions aimed at implementing the desired sequence of state transitions.

Algorithms already exist that can implement such strategies, including the Dijkstra and A* algorithms (*23, 24*) and are commonly used in GPS navigation devices found in cars and cell phones. Many variants of reinforcement learning solve a specific case of this problem, in which the rewarded states are relatively fixed, such as winning a game of Go (*25*). While deep Q networks (*1*) and generative adversarial networks with monte carlo tree search (*25*) are very powerful, what happens when the goals change, or the environmental rules change? In that case, the models may require extensive retraining. The more general problem requires the ability to dynamically recalculate the values associated with each state as circumstances, goals, and set points change, even in novel situations.

Here we explore a computational model that solves this more general problem of how the brain solves problems with changing goals (*26*), and we show how a number of brain regions may implement information processing in ways that correspond to specific model components. The model begins with a core premise: *the brain constitutes a control-theoretic system, generating actions to minimize the discrepancy between actual and desired states*. We developed the Goal-Oriented Learning and Selection of Action (GOLSA) computational neural model from this core premise to simulate how the brain might autonomously learn to solve problems, while maintaining fidelity to known biological mechanisms and constraints such as localist learning laws and real-time neural dynamics. The constraints of biological plausibility both narrow the scope of viable models and afford a direct comparison with neural activity.

The model treats the brain as a high-dimensional control system. It drives behavior to maintain multiple and varying control theoretic set points of the agent’s state, including low level homeostatic (e.g. hunger, thirst) and high level cognitive set points (e.g. a Tower of Hanoi configuration). The model autonomously learns the structure of state transitions, then plans actions to arbitrary goals via a novel hill-climbing algorithm inspired by Dijkstra’s algorithm (*24*). The model provides a domain-general solution to the problem of solving problems and performs well (Supplementary Materials).

The GOLSA model works by representing each possible state of the agent and environment in a network layer, with multiple layers each representing the same sets of states (Figure 1). The Goal Gradient layer is activated by an arbitrarily specified desired (Goal) state and spreads activation backward along possible state transitions represented as edges in the network (*27, 28*). This value spreading activation generates current state values akin to learned state values (Q values) in reinforcement learning, except that the state values can be reassigned and recalculated dynamically as goals change. This additional flexibility allows goals to be specified dynamically and arbitrarily, with all state values being updated immediately to reflect new goals, thus overcoming a limitation of current RL approaches. Essentially, the Goal Gradient is the hill to climb to minimize the discrepancy between actual and desired states in the control theoretic sense. In parallel, regarding the present state of the model system, the Adjacent States layer receives input from a node representing the current state of the agent and environment, which in turn activates representations of all states that can be achieved with one state transition. The valid adjacent states then mask the Goal Gradient layer to yield the Desired Next State representation. In this layer, the most active unit represents a state which, if achieved, will move the agent one step closer to the goal state. This desired next state is then mapped onto an action (i.e. a controller signal) that is likely to effect the desired state transition.

**Figure 1:**
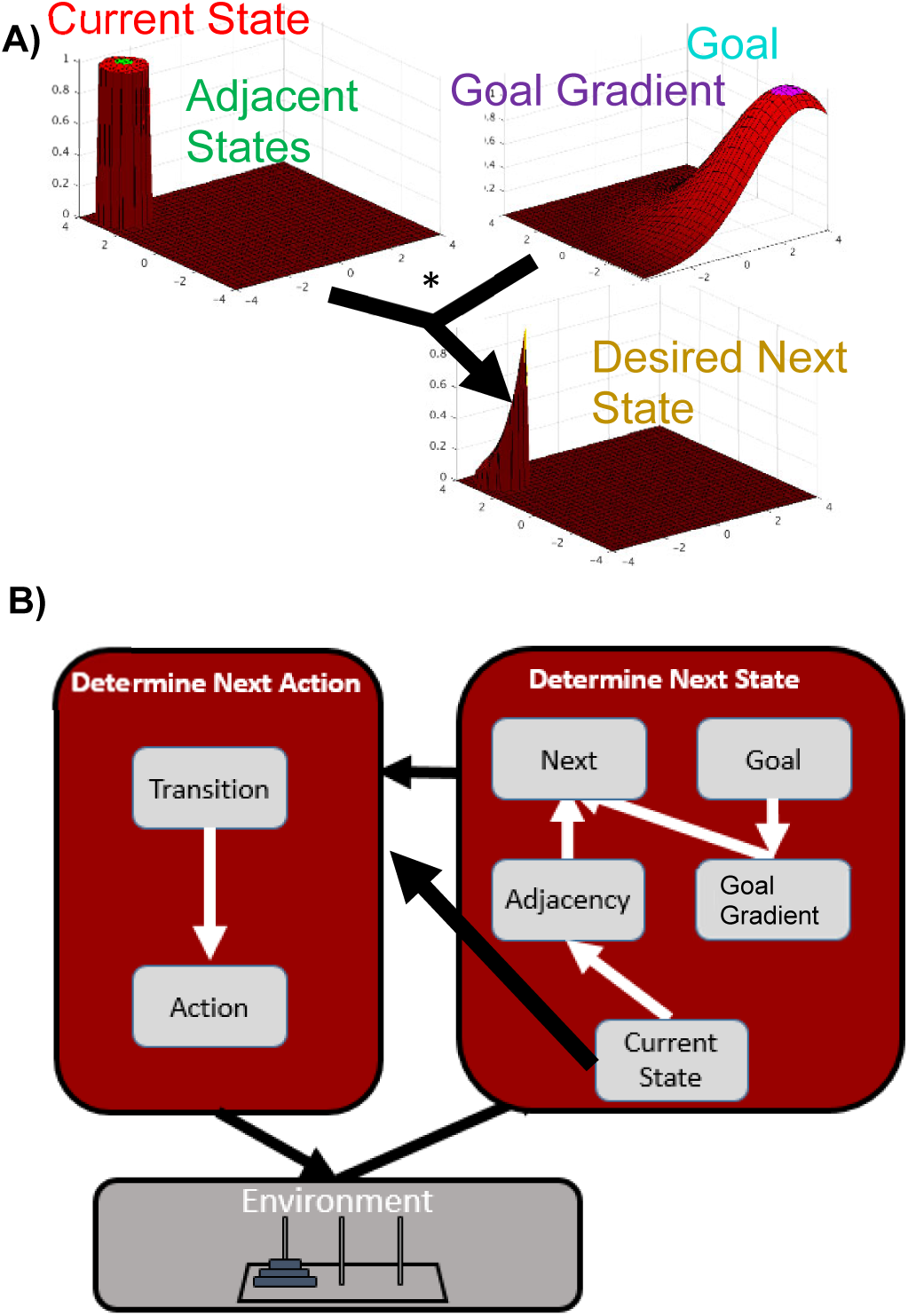
A) The GOLSA model determines the next desired state by hill climbing. Each layer represents the same set of states, one per neuron. Neurons are connected to each other for states that are reachable from another by one action. The Goal state is activated and spreads activation through a Goal Gradient (Proximity) layer, thus dynamically specifying the value of each state given the goal, so that value is greater for states nearer the goal state. The Current State representation activates all Adjacent States, i.e. that can be achieved with one state transition. These adjacent states mask the Goal Gradient input to the Desired Next State, so that the most active unit in the Desired Next State represents a state attainable with one state transition and which will bring the state most directly toward the goal state. The font colors match the model layer to corresponding brain regions in Figs. 3 and 4. **B) The desired state transition** is determined by the conjunction of current state and desired next state. The GOLSA model learns a mapping from desired state transitions to the actions that cause those transitions. After training, the model can generate novel action sequences to achieve arbitrary goal states. Adapted from (*26*).

The details of the model implementation are in the Supplementary materials, and the model code is available online (Supplementary Materials). Behaviorally, we found that the GOLSA model is able to learn to solve arbitrary problems, such as reaching novel states in the Tower of Hanoi task (Figure 2A). It does this without hard-wired knowledge, simply by making initially random actions and learning from the outcomes, then synthesizing the learned information to achieve whatever state is specified as the goal state.

**Figure 2:**
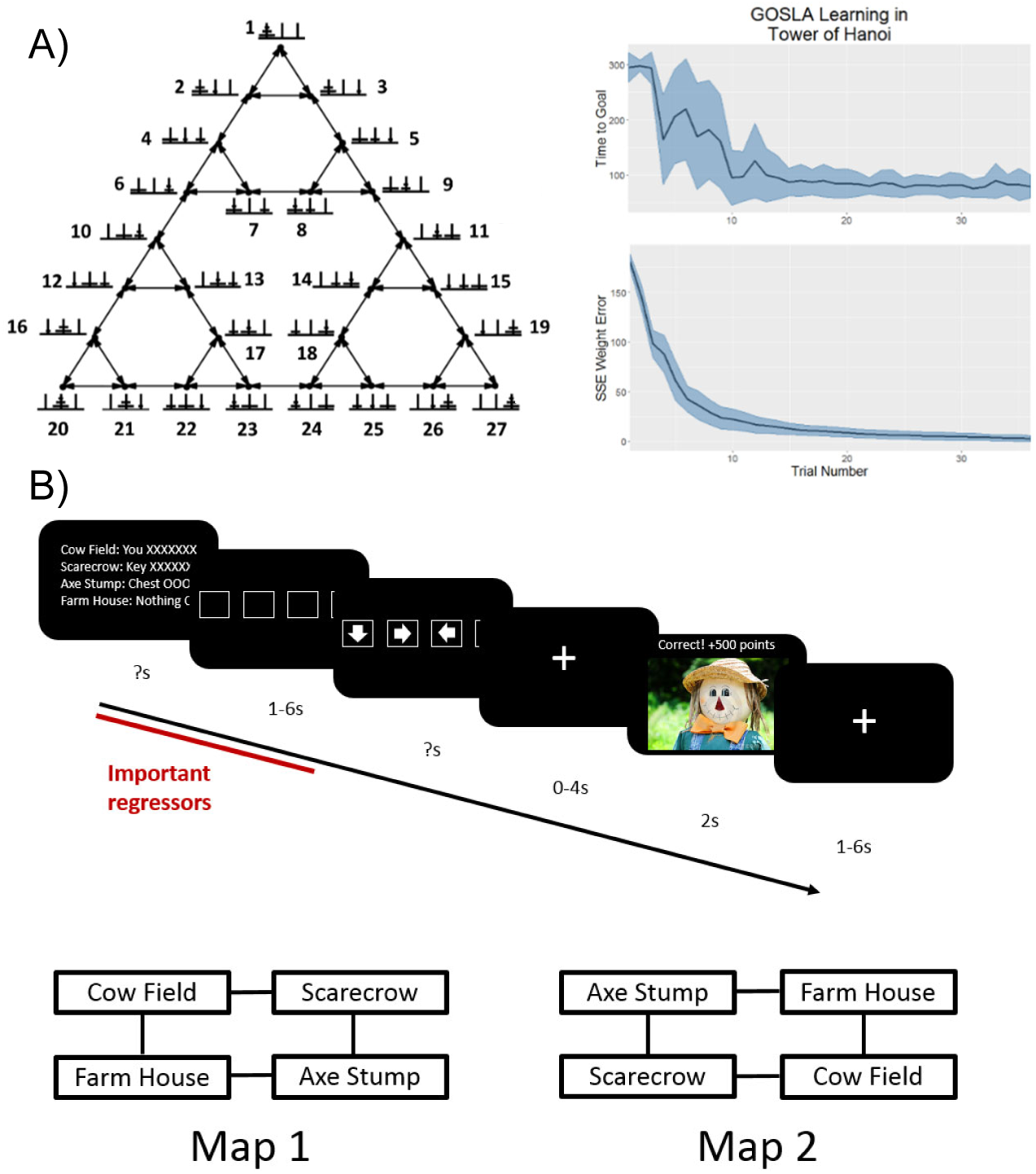
A) The GOLSA model learns to solve problems, achieving arbitrary goal states. It does this by making arbitrary actions and observing which actions cause which state transitions. Figure adapted from (*26, 29*). **B) Treasure Hunt task** Both the GOLSA model and the human fMRI subjects performed a simple treasure hunt task, in which subjects were placed in one of four possible starting locations, then asked to generate actions to reach any of the other possible locations. To test multi-step transitions, subjects had to first move to the location of a key needed to unlock a treasure chest, then move to the treasure chest location. The mapping of finger buttons to game movements was random on each trial and revealed after subjects were given the task and had to plan their movements, thus avoiding motor confounds during planning.

Having found that the model can learn autonomously to solve arbitrary problems, we then aimed to identify which brain regions might show representations and activity that matched particular GOLSA model layers. To do this, we tested the GOLSA model with a Treasure Hunt task (Figure 2B and Supplementary Materials), which was performed by both the GOLSA model and human subjects with fMRI. Subjects were placed in one of four starting states and had to traverse one or two states to achieve a goal, by retrieving a key and subsequently using it to unlock a treasure chest for a reward (Figure 2B).

To analyze the fMRI and model data, we used model-based fMRI with representational similarity analysis (RSA) (*30*) (Supplementary Materials). Briefly, each representational dissimilarity matrix (RDM) represented the pairwise correlations across 96 total patterns – 4 starting states by 8 trial types by 3 time points within a trial (problem description, response, and feedback). For each model layer, the pairwise correlations are calculated with the activity pattern across layer cells in one condition vs. the activity pattern in the same layer in the other condition. For each voxel in the brain, the pairwise correlations are calculated with the activity pattern in a local neighborhood of voxels around the voxel in question, for one condition vs. the other condition. The comparison between GOLSA model and fMRI RDMs consists of looking for positive correlations between elements of the upper symmetric part of a given GOLSA model layer RDM vs. the RDM around a given voxel in the fMRI RDMs. The resulting fMRI RSA maps, one per GOLSA model layer, show which brain regions have representational similarities between particular model components and particular brain regions. The fMRI RSA maps are computed for each subject and then tested across subjects for statistical significance in a given brain region. Full results are in Table S3.

We found that the patterns of activity in a number of distinct brain regions match those expected of a control theoretic system, as instantiated in the GOLSA model (Figures 3, 4). Orbitofrontal cortex (OFC) activity patterns match model components that represent both a cognitive map (*6*) and a flexible goal value representation (*7*), specifically matching the Goal and Goal Gradient layer activities. These layers represent the current values of the goal state and the current values of states near the goal state, respectively. The Goal Gradient layer incorporates cognitive map information in terms of which states can be reached from which other states. This suggests mechanisms by which OFC regions may calculate the values of states dynamically as part of a value-based decision process, by spreading activation of value from a currently active goal state representation backward. The GOLSA model representations of the desired next state also match overlapping regions in the orbitofrontal cortex (OFC) and ventromedial prefrontal cortex (vmPFC), consistent with a role in finding the more valuable decision option (Figure 3). Reversal learning and satiety effects as supported by the OFC reduce to selecting a new goal state or deactivating a goal state respectively, which immediately updates the values of all states. Collectively this provides a mechanistic account of how value-based decision-making functions in OFC and vmPFC.

**Figure 3:**
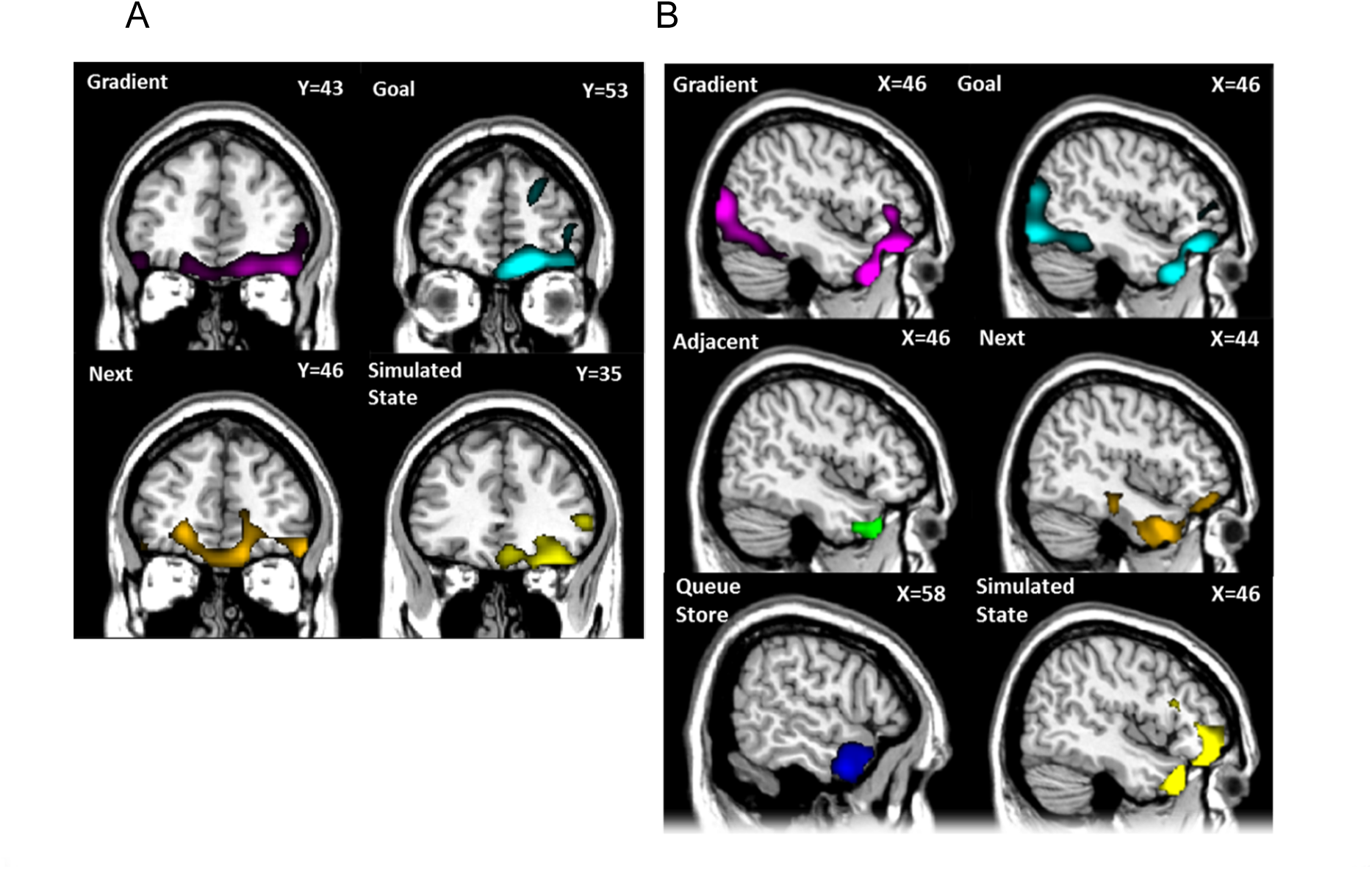
Representational Similarity Analysis (RSA) of model layers vs. human subjects performing the same Treasure Hunt task. All results shown are significant clusters across the population with a cluster defining threshold of p < 0.001 cluster corrected to p < 0.05 overall. **(A)** population Z maps showing significant regions of similarity to model layers in orbitofrontal cortex. Cf. Figure 1 and Figure S1B. The peak regions of similarity for goal-gradient and goal show considerable overlap in right OFC. The region of peak similarity for simulated-state is more posterior. To most clearly show peaks of model-image correspondence, the maps of gradient and goal are here visualized at p < 0.00001 while all others are visualized at p < 0.001. **(B)** Z maps showing significant regions of similarity to model layers in right temporal cortex. The peak regions of similarity for goal-gradient and goal overlap and extend into the OFC. The peak regions of similarity for adjacent-states, next-desired-state, and -simulated-state occur in similar but not completely overlapping regions, while the cluster for queue-store is more lateral.

**Figure 4:**
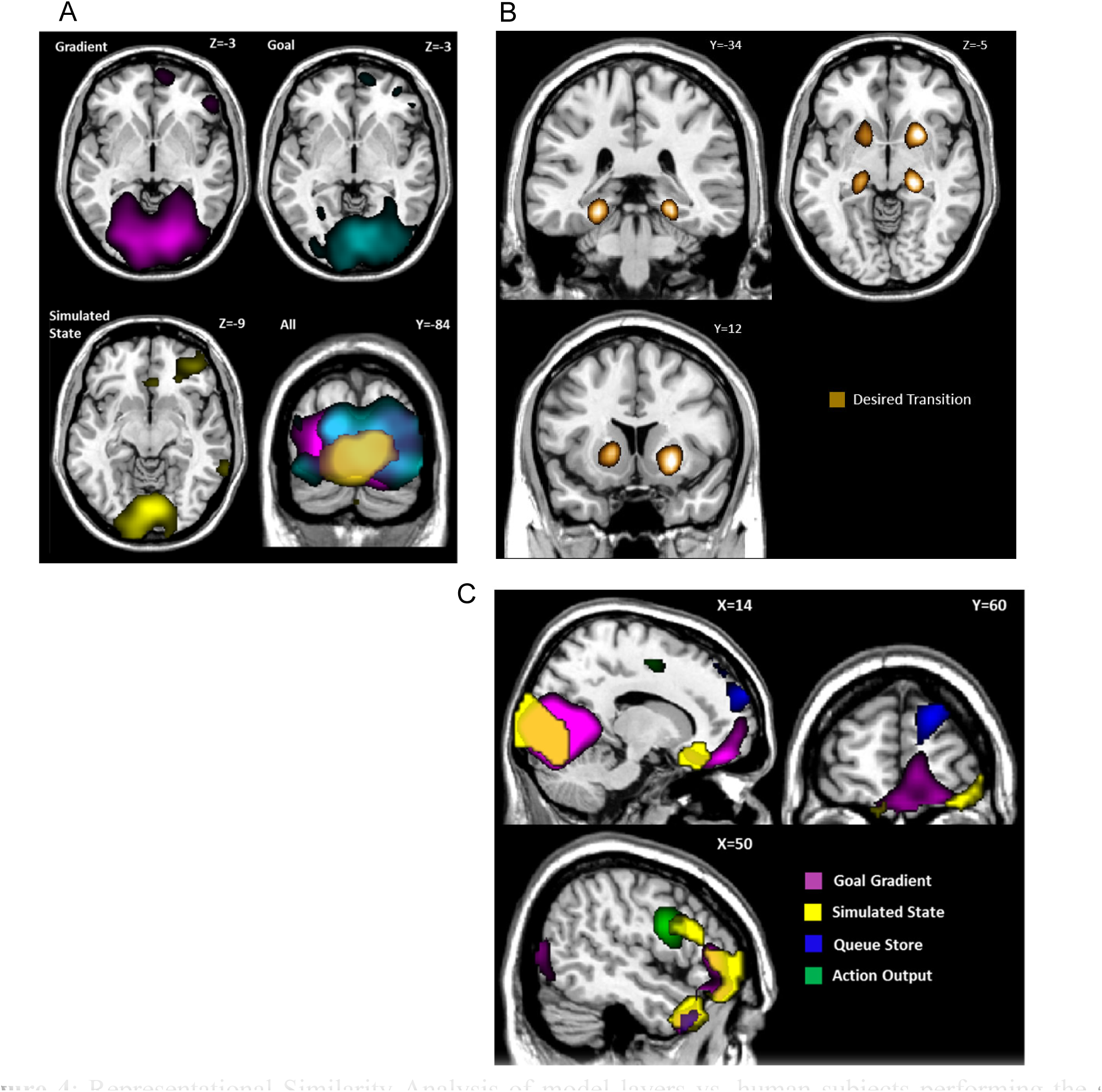
Representational Similarity Analysis of model layers vs. human subjects performing the same Treasure Hunt task, with the same conditions as in Figure 3. **(A)** Population Z maps showing significant regions of similarity to model layers in visual cortex. The peak regions of similarity for goal-gradient and goal overlap substantially, primarily in bilateral cuneus, inferior occipital gyrus, and lingual gyrus. The simulated-state layer displayed significantly similar activity to that in a smaller medial and posterior region. Statistical thresholding and significance are the same as Figure 3. **(B)** Z map showing significant regions of similarity to the desired-transition layer. Similarity peaks were observed for desired-transition in bilateral hippocampal gyrus as well as bilateral caudate and putamen. The desired-transition map displayed here was visualized at p < 0.00001 for clarity. **(C)** Z maps showing significant regions of similarity to the model layers in frontal cortex. Similarity peaks were observed for queue-store in superior frontal gyrus (BA10). Action-output activity most closely resembled activity in inferior frontal gyrus (BA9), while simulated-state and goal-gradient patterns of activity were more anterior (primarily BA45). Similarity between activity in the latter two layers and activity in OFC, visual cortex, and temporal pole is also visible.

The GOLSA model also incorporates a mechanism that allows multi-step planning, by representing a Simulated State as if the desired next state were already achieved, so that the model can plan multiple subsequent state transitions iteratively prior to committing to a particular course of action (Figure S1B). Those subsequent state transitions are represented in a Queue Store layer pending execution via competitive queueing. This constitutes a mechanism of prospection (*31*) and planning (*32*). The Simulated State layer in the GOLSA model shows strong representational similarity with regions of the OFC and anterior temporal lobe, and the Queue Store layer shows strong similarity with the anterior temporal lobe and lateral prefrontal cortex. This constitutes a mechanistic account of how the vmPFC and OFC in particular might contribute to multi-step goal-directed planning, and how plans may be stored in lateral prefrontal cortex.

The visual cortex also shows representational patterns consistent with representing the goal, goal gradient, and simulated future states (Figures 3B and 4). This suggests a role for the visual cortex in planning, in the sense of representing anticipated future states beyond simply representing current visual input. One possibility is that this reflects an attentional effect that facilitates processing of visual cues representing anticipated future states. Another possibility is that visual cortex represents a kind of working memory for anticipated future visual states (*13*). In either case, the results are consistent with the notion that the visual cortex may not be only a sensory region but may play some role in planning by representing the details of anticipated future states.

The anterior temporal lobe likewise shows representations of the goal, goal gradient, the adjacent states, the next desired state, and simulated future and queue store states (Figures 3B, 4C). In one sense this is not surprising, as the states of the task are represented by images of objects, and visual objects (especially faces) are represented in the anterior temporal lobe (*33*). Still, the fact that the anterior temporal lobe shows representations consistent with planning mechanisms suggests a more active role in planning beyond feedforward sensory processing as commonly understood (*34*).

Once the desired next state is specified, it must be translated to an action. The hippocampus and striatum match the representations of the Desired Transition layer in the GOLSA model. This model layer represents a conjunction of the current state and desired next state transitions, which in the GOLSA model is a necessary step toward selecting an appropriate action to achieve the desired transition. This is consistent with the role of the hippocampus in prospection (*31*), and it suggests computational and neural mechanisms by which the hippocampus may play a key role in turning goals into predictions about the future, for the purpose of planning actions (*9, 10*). Finally, as would be expected, the motor output representations in the GOLSA model match motor output patterns in the motor cortex (Figure 4C).

The results above show how a computational neural model, the GOLSA model, provides a novel computational account of a number of brain regions. The guiding theory is that a substantial set of brain regions function together as a control-theoretic mechanism, generating behaviors to minimize the discrepancy between the current state and the desired (goal) state.

Our findings bear a resemblance to the Free Energy principle. According to this, organisms learn to generate predictions of the most likely (i.e. rewarding) future states under a policy, then via active inference emit actions to cause the most probable outcomes to become reality, thus minimizing surprise (*35, 36*). Like active inference, the GOLSA model emits actions to minimize the discrepancy between the actual and predicted state. Of note, the GOLSA model specifies the future state as a desired state rather than a most likely state. This allows a state that has a high current value to be pursued, even if the probability of being in that state is very low (for example buying lottery tickets). Furthermore, the model includes the mechanisms of Figure 1, which allow for flexible planning given arbitrary goals. The GOLSA model is a process model and simulates neural activity as a dynamical system, which affords the more direct comparison with neural activity representations over time as in Figures 3 and 4.

The GOLSA model shares some similarity with model-based reinforcement learning (MBRL), in that both include learned models of next-state probabilities as a function of current state and action pairs. Still, a significant limitation of both model-based and model free RL is that typically there is only a single ultimate goal, e.g. gaining a reward or winning a game. Q-values (*37*) are thus learned in order to maximize a single reward value. This implies several limitations: (1) that Q values are strongly paired with corresponding states; (2) that there is only one Q value per state at a given time, as in a Markov decision process (MDP), and (3) Q values are generally updated via substantial relearning. In contrast, real organisms will find differing reward values associated with different goals at different times and circumstances. This implies that goals will change over time, and re-learning Q-values with each goal change would be inefficient. Instead, a more flexible mechanism will dynamically assign values to various goals and then plan accordingly. The GOLSA model exemplifies this approach, essentially replacing the learned Q values of MBRL and MDPs with an activation-based representation of state value, which can be dynamically reconfigured as goals change. This overcomes the three limitations above.

The GOLSA model may in principle be extended hierarchically. The frontal cortex has a hierarchical representational structure, in which higher levels of a task may be represented more anteriorly (*38*). Such hierarchical structure has been construed to represent higher, more abstract task rules (*12, 39*). The GOLSA model suggests another perspective, that higher level representations consist of higher level goals instead of higher level rules. In the coffee-making task for example (*40*), the higher level task of making coffee may require a lower level task of boiling water. If the GOLSA model framework were extended hierarchically, the high level goal of having coffee prepared would activate a lower level goal of having the water heated to a specified temperature. The goal specification framework here is intrinsically more robust than a rule or schema based framework – rules may fail to produce a desired outcome, but if an error occurs in the GOLSA task performance, replanning simply calculates the optimal sequence of events from whatever the current state is, and the error will be automatically addressed.

The main limitation of the GOLSA model is its present implementation of one-hot state representations. This makes a scale-up to larger and continuous state spaces challenging. Future work may overcome this limitation by replacing the one-hot representations with vector-valued state representations and the learned connections with deep network function approximators. This would require corresponding changes in the search mechanisms of Figure 1A, from parallel, spreading activation to a serial, monte carlo tree search mechanism. This would be consistent with evidence of serial search during planning (*41, 42*) and would afford a new approach to artificial general intelligence that is both powerful and similar to human brain function.

## Acknowledgments

We thank A. Ramamoorthy for helpful discussions and J. Fine and W. Alexander for helpful comments on the manuscript. Supported by the Indiana University Imaging Research Facility. JWB was supported by NIH R21 DA040773.

## Author contributions

JWB and NZ designed the model and experiment. NZ implemented and simulated the model, implemented and ran the fMRI experiment, and analyzed the data. JWB and NZ wrote the paper.

