## Supplementary material for "Foundations of human problem solving"

N. Zarr[1], J.W. Brown[1]*.

**This PDF file includes:**

Materials and Methods
Supplementary Text
Figs. S1 to S4
Tables S1 to S3

### Materials and Methods

### Model Components

#### Layers

The GOLSA model is constructed from a small set of basic components, and the model code is freely available as supplementary material. The main component class is a layer of units, where each unit represents a neuron (or, more abstractly, a small subpopulation of neurons) corresponding to either a state, a state transition, or an action. The activity of units in a layer represents the neural firing rate and is instantiated as a vector updated according to a first order differential equation (c.f. Grossberg, 1973). The activation function varies between layers, but all units in a particular layer are governed by the same equation. The most typical activation function for a single unit is,

$$da(t) = \left(-\lambda a(t)dt + \left(1 - a(t)\right)Edt - Idt + \varepsilon N(t)\sqrt{dt}\right), \qquad \text{(S1)}$$

where *a* represents activation, i.e. the firing rate, of a model neuron. The five terms of this equation represent passive decay, shunting excitation, linear inhibition, and random noise. “Shunting” refers to the fact that excitation (E) scales inversely as current activity increases, with a natural upper bound of 1. The passive decay works in a similar fashion, providing a natural lower bound activity of 0. The inhibition term linearly suppresses unit activity, while the final term adds normally distributed noise ($\mu$=0, $\sigma$=1) . Because the differential equations are approximated using the Euler method, the noise term is multiplied by $\sqrt{dt}$ to standardize the magnitude across different choices of dt (Busemeyer & Townsend, 1993; Usher & McClelland, 2001). The speed of activity change is determined by a time constant $\tau$. The parameters $\tau$, $\lambda$, $\varepsilon$ vary by layer in order to implement different processes. E and I are the total excitation and inhibition impinging on a particular unit for every presynaptic unit j in every projection p onto the target unit,

$$E = \sum_{p} \sum_{j} [w_{p_j} a_{p_j}]^{+} \quad \text{(S2)}$$

$$I = \sum_{p} \sum_{j} [w_{p_j} a_{p_j}]^{-} \quad \text{(S3)}$$

A second activation function used in several places throughout the model is,

$$da(t) = \left(-\lambda a(t)dt + \left(1 - a(t)\right)Edt - a(t)Idt + \varepsilon N(t)\sqrt{dt}\right) \quad \text{(S4)}$$

This function is identical to Equation () except that the inhibition is also shunting, such that it exhibits a strong effect on highly active units and a smaller effect as unit activity approaches 0. While more typical in other models, shunting inhibition has a number of drawbacks in the current model. Two common uses for inhibition in the GOLSA model are winner-take-all dynamics and regulatory inhibition which resets layer activity. Shunting inhibition impedes both of these processes because inhibition fails to fully suppress the appropriate units, since it becomes less effective as unit activity decreases.

**Projections**

Layers connect to each other via projections, representing the synapses connecting one neural population to another. The primary component of projections is a weight matrix specifying the strength of connections between each pair of units. Learning is instantiated by updating the weights according to a learning function. These functions vary between the projections responsible for the model learning and are fully described in the section below dealing with each learning type. Some projections also maintain a matrix of traces updated by a projection-specific function of presynaptic or postsynaptic activity. The traces serve as a kind of short-term memory for which pre or postsynaptic units were recently activated, which serve a very similar role to eligibility traces as in Barto et al. (1979), though with a different mathematical form.

**Nodes**

Nodes are model components that are not represented neurally via an activation function. They represent important control and timing signals to the model and are either set externally or update autonomously according to a function of time. For instance, sinusoidal oscillations are used to gate activity between various layers. While in principle rate-coded model neurons could implement a sinusoidal wave, the function is simply hard coded into the update function of the node for simplicity. In some cases, it is necessary for an entire layer to be strongly inhibited when particular conditions hold true, such as when an oscillatory node is in a particular phase. Layers therefore also have a list of inhibitor nodes that prevent unit activity within the layer when the node value meets certain conditions. In a similar fashion, some projections are gated by nodes such that they allow activity to pass through and/or allow the weights to be updated only when the relevant node activity satisfies a particular condition. Another important node provides strong inhibition to many layers when the agent changes states.

**Environment**

The agent operates in an environment consisting of discrete states, with a set of allowable state transitions. Allowable state transitions are not necessarily bidirectional, but for the present simulations, they are deterministic (unlike the typical MDP formulation used in RL). In some simulations, the environment also contains different types of reward located in various states, which can be used to drive goal selection. In other simulations, the goal is specified externally via a node value.

#### Complete Network

Each component and subnetwork of the model is described in detail below or in the main text, but for reference and completeness a full diagram of the core network is shown in Figure S1A, and the network augmented for multi-step planning is shown in Figure S1B. Some of the basic layer properties are summarized in Table S1. Layers and nodes are referred to using italics, such that the layer representing the current state is referred to simply as *current-state*.

#### Representational structure

In Figure S1B, the layers Goal, Goal Gradient, Next State, Adjacent States, Previous States, Simulated State, and Current State all have the same number of nodes and the same representational structure, i.e. one state per node.

The layers Desired Transition, Observed Transition, Transition Output, Queue Input, Queue Output, and Queue Store likewise have the same representational structure, which is the number of possible states squared. This allows a node in these layers to represent a transition from one specific state to another specific state.

The layers Action Input, Action Output, and Previous Action all have the same representational structure, which is one possible action per node.

A)

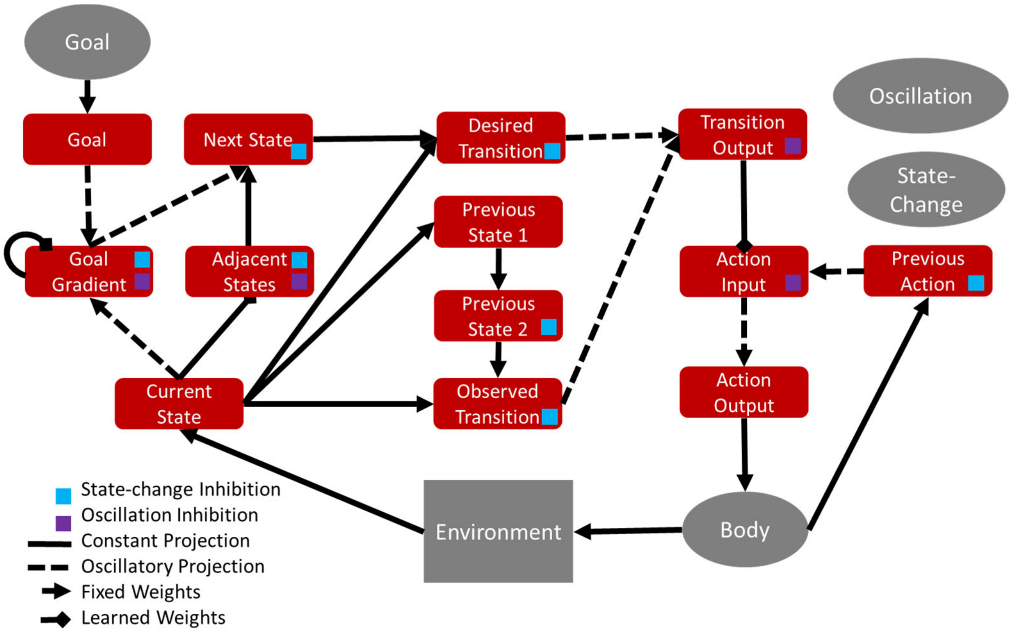


B)

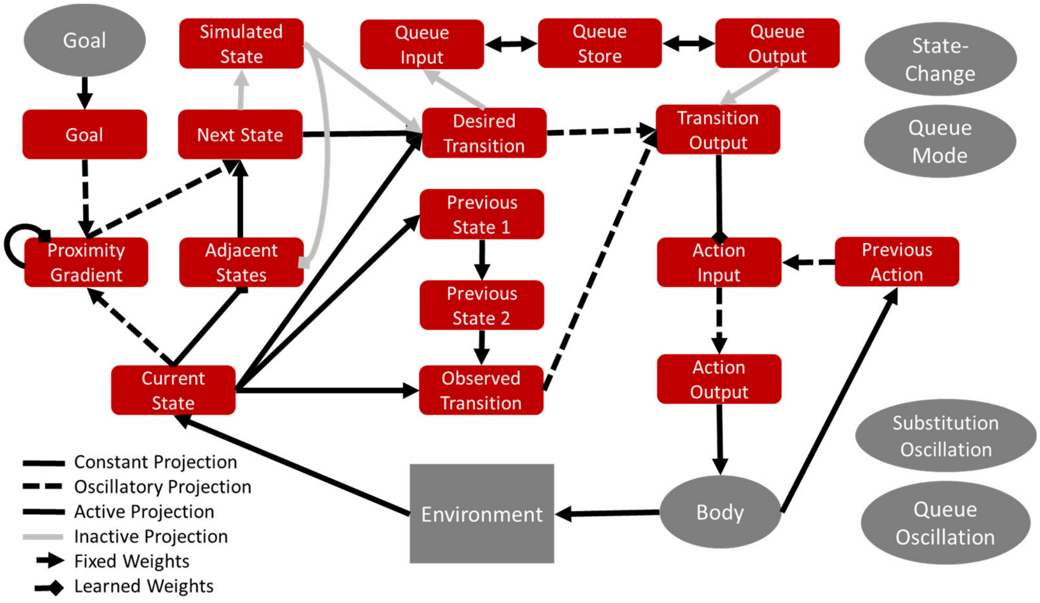


**Figure S1.** (A) Full diagram of core model. Each rectangle represents a layer and each arrow a projection. The body is a node, and two additional nodes are not shown which provide inhibition at each state-change and oscillatory control. The colored squares indicate which layers receive inhibition from these nodes. Some recurrent connections not shown. (B) Full diagram of extended model, with added top row representing ability to plan multiple state transition steps ahead (Simulated State, Queue Input, Queue Store, and Queue Output layers).

| Layer Name | Inhibition Type | Time Constant | Decay Rate | Noise Gain |
|---|---|---|---|---|
| current-state | shunting | 0.5 | 1 | 0 |
| goal | linear | 1 | 1 | 0 |
| goal-gradient | linear | 1 | 1 | 0 |
| adjacent-states | linear | 0.2 | 1 | 0.01 |
| next-desired-state | linear | 1 | 0.1 | 0 |
| previous-state-1 | shunting | 4 | 0.5 | 0 |
| previous-state-2 | linear | 1 | 0.001 | 0 |
| previous-action | linear | 1 | 0.001 | 0 |
| observed-transition | linear | 1.5 | 1 | 0 |
| desired-transition | linear | 1.5 | 1 | 0 |
| transition-output | linear | 1.5 | 1 | 0 |
| action-input | linear | 1 | 1 | 0 |
| action-output | linear | 0.5 | 0.2 | 0 |

**Table S1. Full list of core model layers and associated parameters.**

### Task description

The goal-pursuit task (Figure S2) was created and presented in OpenSesame, a Python-based toolbox for psychological task design (Mathôt, Schreij, & Theeuwes, 2012). In the task, participants control an agent which can move within a small environment comprised of four distinct states. The nominal setting is a farm, and the states are a field with a scarecrow, the lawn in front of the farm house, a stump with an axe, and a pasture with cows. Each is associated with a picture of the scene obtained from the internet[1]. These states were chosen to exemplify categories previously shown to elicit a univariate response in different brain regions, namely faces, houses, tools, and animals (Anzellotti, Mahon, Schwarzbach, & Caramazza, 2011; Kanwisher, McDermott, & Chun, 1997; O'Craven, Downing, & Kanwisher, 1999).

Over the course of the experiment, participants were told the locations of treasure chests and the keys needed to open them. By arriving at a chest with the key, participants earned points which were converted to a monetary bonus at the end of the experiment. The states were arranged in a square, where each state was accessible from the two adjacent states but not the state in the opposite corner (diagonal movement was not allowed).

Each trial began with the presentation of a text screen displaying the relevant information for the next trial, namely the locations of the participant, the key, and the chest (Figure S2). Because the neural patterns elicited during the presentation were the primary target of the decoding analysis, it was important that visual information be as similar as possible across different goal configurations, to avoid potential confounds. To hold luminance as constant as possible across conditions, each line always had the same number of characters. Since, for instance, "Farm House: key" has fewer characters than "Farm House: Nothing", filler characters

[1] All images were labeled as available for reuse

were added to the shorter lines, namely Xs and Os. On some trials Xs were the filler characters on the top row and Os were the filler characters on the bottom rows. This manipulation allowed us to attempt to decode the relative position of the Xs and Os to test whether decoding could be achieved due only to character-level differences in the display. We found no evidence that our results reflect low level visual confounds such as the properties of the filler characters.

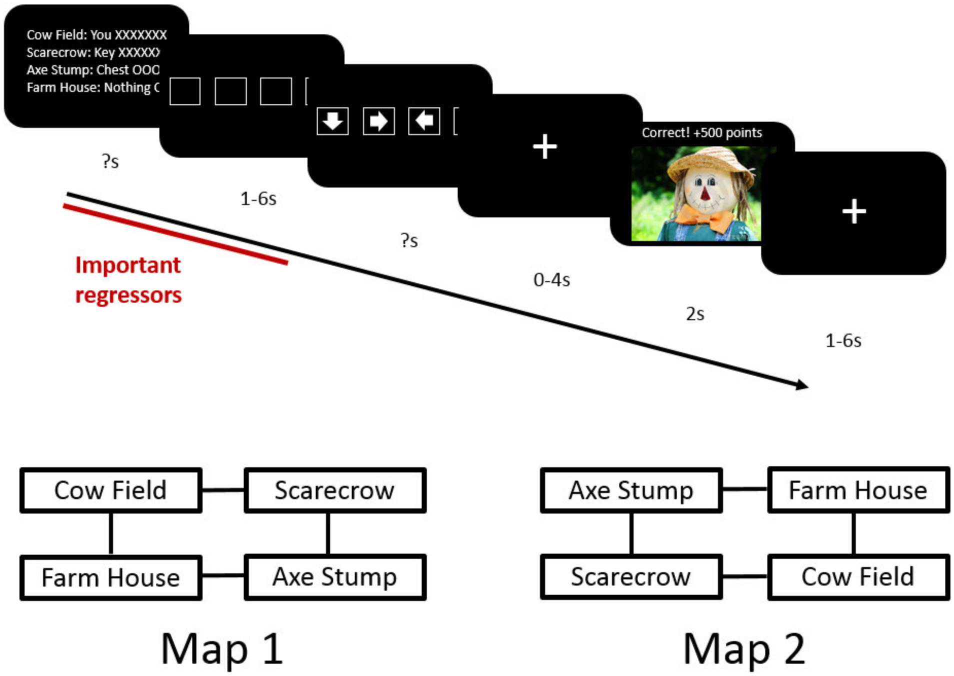


**Figure S2.** Top: Behavioral task sequence. Participants first saw an information screen specifying the contents of each of the four states ('you', 'key', 'chest', or 'nothing'). After a jittered delay, participants selected a desired movement direction and after another delay saw an image of the outcome location. A '?' indicates a self-paced response. Bottom: The two state-space maps used in the experiment. One map was used in the first half of trials while the other was used in the second half, in counterbalanced order.

Participants were under no time constraint on the information screen and pressed a button when they were ready to continue. A delay screen then appeared consisting of four empty boxes. After a jittered interval (1-6s, distributed exponentially), arrows appeared in the boxes. The arrows represented movement directions and the boxes corresponded to four buttons under the participants left middle finger, left index finger, right index finger, and right middle finger, from left to right. Participants pressed the button corresponding to the box with the arrow pointing in the desired direction to initiate a movement. A fixation cross then appeared for another jittered delay of 0-4s, followed by a 2s display of the newly reached location if their choice was correct or an error screen if it was incorrect.

If the participant did not yet have the key required to open the chest, the correct movement was always to the key. Sometimes the key and chest were in the same location in which case the participant would earn points immediately. If they were in different locations, then on the next trial the participant had to move to the chest. This structure facilitated a mix of goal distances (one and two states away) while controlling the route required to navigate to the goal.

If the chosen direction was incorrect, participants saw an error screen displaying text and a map of the environment. Participants advanced from this screen with a button press and then restarted the failed trial. If the failed trial was the second step in a two-step sequence (i.e., if they had already gotten the key and then moved to the wrong state to get to the chest), they had to repeat the previous two trials.

Repeating the failed trial ensured that there were balanced numbers of each class of event for decoding, since an incorrect response indicated that some information was not properly maintained or utilized. For example, if a participant failed the second step of a two-trial sequence, then they may not have properly encoded the final goal when first presented with the information screen on the previous trial, which specified the location of the key and the chest.

Halfway through the experiment, the map was reconfigured such that states were swapped across the diagonal axes of the map. This was necessary because otherwise, each state could be reached by exactly two movement directions and exactly two movement directions could be made from it. For instance, if the farm house was the state in the lower left, the farmhouse could only be reached by moving left or down from adjacent states, and participants starting at the farm house could only move up or to the right. If this were true across the entire experiment, above-chance classification of target state, for instance, could appear in regions that in fact only contain information about the intended movement direction.

Each state was the starting state for one quarter of the trials and the target destination for a different quarter of the trials. All trials were one of three types. One category consisted of single-trial (*single*) sequences in which the chest and key were in the same location. The sequences in which the chest and key were in separate locations required two trials to complete, one to move from the initial starting location to the key and another to move from the key location to the chest location. These two steps formed the other two classes of trials, the first-of-two (*first*) and second-of-two *(second*) trials. Recall that on *second* trials, no information other than the participant's current location is presented on the starting screen to ensure that the participant maintained the location of the chest in memory across the entire two-trial sequence (if it was presented on the *second* trial, there would be no need to maintain that information through the *first* trial). The trials were evenly divided into *single*, *first*, and *second* classes with 64 trials in each class. Therefore, every trial had a starting state and an immediate goal, while one third of trials also had a more distant final goal.

Immediately prior to participating in the fMRI version of the task, participants completed a short 16-trial practice outside the scanner to refresh their memory. Before beginning the first run inside the scanner, participants saw a map of the farm states and indicated when they had memorized it before moving on. Within each run, participants completed as many trials as they could within eight minutes. As described above, exactly halfway through the trials, the state space was rearranged with each state moving to the opposite corner. Therefore, when participants completed the first half of the experiment, the current run was terminated and participants were given time to learn the new state space before scanning resumed. At the end of the experiment, participants filled out a short survey about their strategy.

**Participants**

In total, 49 participants (28 female) completed the behavioral-only portion of the experiment, including during task piloting (early versions of the behavioral task were slightly different than described below). Participants provided written informed consent in accordance with the Institutional Review Board at Indiana University, and were compensated $10/hour for their time plus a performance bonus based on accuracy up to an additional $10. The behavioral task first served as a pilot during task design and then as a pre-screen for the fMRI portion, in that only participants with at least 90% accuracy were invited to participate. Additional criteria for scanning were that the subjects be right handed, free of metal implants, free of claustrophobia, weigh less than 440 pounds, and not be currently taking psychoactive medication. In total, 25 participants participated in the fMRI task but one subject withdrew shortly after beginning, leaving 24 subjects who completed the imaging task (14 female).

**fMRI acquisition and data preprocessing**

Imaging data were collected on a Siemens Magnetom Trio 3.0-Tesla MRI scanner and a 32 channel head coil. Foam padding was inserted around the sides of the head to increase participant comfort and reduce head motion. Functional T2* weighted images were acquired using a multiband EPI sequence (Moeller et al., 2010) with 42 contiguous slices and 3.44 x 3.44 x 3.4 $mm^3$ voxels (echo time = 28 ms; flip angle = 60; field of view = 220, multiband acceleration factor = 3). For the first subject, the TR was 813ms, but during data collection for the second subject the TR changed to 816ms for unknown reasons. The scanner was upgraded after collecting data from an additional five subjects, at which point the TR remained constant at 832 ms. All other parameters remained unchanged. High-resolution $T_1$ – weighted MPRAGE images were collected for spatial normalization (256 x 256 x 160 matrix of 1 x 1 x 1$mm^3$ voxels, TR = 1800, echo time = 2.56 ms; flip angle = 9).

Functional data were spike-corrected using AFNI's 3dDespike (http://afni.nimh.nih.gov/afni). Functional images were corrected for difference in slice timing using sinc-interpolation and head movement using a least-squares approach with a 6-parameter rigid body spatial transformation. For subjects who moved more than 3mm total or 0.5 mm between TRs, 24 motion regressors were added to subsequent GLM analyses.

Because MVPA and representation similarity analysis (RSA) rely on precise voxelwise patterns, these analyses were performed before spatial normalization. For the univariate analyses, structural data were coregistered to the functional data and segmented into gray and white matter probability maps (Ashburner & Friston, 1997). These segmented images were used to calculate spatial normalization parameters to the MNI template, which were subsequently applied to the functional data. As part of spatial normalization, the data were resampled to 2 x 2 x 2$mm^3$. An 8-

mm full-width/half-maximum isotropic Gaussian smoothing was applied to the functional images. All analyses included a temporal high-pass filter (128s) and correction for temporal autocorrelation using an autoregressive AR(1) model.

**Univariate GLM**

For initial univariate analyses, we measured the neural response associated with each outcome state at the outcome screen (when an image of the state was displayed), as well as the signal at the start of the trial associated with each immediate goal location. Five timepoints were modeled in the GLM used in this analysis, namely the start of the trial, the button press to advance, the appearance of the arrows and subsequent response, the start of the feedback, and the end of the feedback. The regressors marking the start of the trial and the start of the feedback screen were further individuated by the immediate goal on the trial. A separate error regressor was used when the response was incorrect, meaning they did not properly pursue the immediate goal and received error feedback. All correct trials in which participants moved to, for instance, the cow field, used the same trial start and feedback start regressors.

The GLM was fit to the normalized and smoothed functional images. The resulting beta maps were combined at the second level with a voxel-wise threshold of $p < 0.001$ and cluster corrected ($p < 0.05$) to control for multiple comparisons. We assessed the univariate response associated with each outcome location, by contrasting each particular outcome location with all other outcome locations. The response to the error feedback screen was assessed in a separate contrast against all correct outcomes. To test for any univariate responses related to the immediate goal, we performed an analogous analysis using the trial start regressors which were individuated based on the immediate goal. For example, the regressor 'trialStartHouseNext' was associated with the beginning of every trial where the farmhouse was the immediate goal location. To assess the univariate signal associated with the farmhouse immediate goal, we performed a contrast between this regressor and all other trial start regressors.

**Representational similarity analysis (RSA)**

As before, a GLM was fit to the realigned functional images. The following events were modeled with impulse regressors: trial onset (information screen), key press to advance to the decision screen, the prompt and immediately subsequent action (modeled as a single regressor), the onset of the outcome screen, and the termination of the outcome screen. The RSA analysis used beta maps derived from the regressors marking trial onset, prompt/response, and outcome screen onset.

Each of these regressors (except those used in error trials) were further individuated by the (start state, next state, final goal) triple constituting the goal path. There were 8 distinct trial types starting in each state. Each state could serve as the starting point of two single-step sequences (in which the key and treasure chest are in the same location) and four two-step sequences (in which the key and treasure chest are in different locations). Each state could also be the midpoint of a two-step sequence with the treasure chest located in one of two adjacent states. With three regressors used for each trial, there were 4 starting states * 8 trial types * 3 time points = 96 total patterns used to create the Representational Dissimilarity Matrix (RDM) in each searchlight region, where cell $x_{ij}$ in the RDM is defined as one minus the Pearson correlation between the $i^{th}$ and $j^{th}$ patterns. Values close to 2 therefore represent negative correlation (high representational distance) while values close to 0 indicate a positive correlation (low representational distance).

**Model RDMs**

To derive the model-based RDMs, the GOLSA model was run on an analogue of the goal pursuit task, using a four state state-space with four actions corresponding to movement in each cardinal direction. The model layer timecourses of activity are shown in Figures S3 and S4 for one- and two-step trials, respectively. The base GOLSA model is not capable of maintaining a plan across an arbitrary delay, but instead acts immediately to make the necessary state transitions. The competitive queue (Bullock, 2004) module allows state transition sequences to be maintained and executed after a delay, and was therefore necessary to model the task in the most accurate manner possible. However, the goal-learning module was not necessary since goals were externally imposed. Because participants had to demonstrate high performance on the task before entering the scanner, little if any learning took place during the experiment. As a result, the model was trained extensively on the state space before performing any trials used in data collection. To further simulate likely patterns of activity in the absence of significant learning, the input from *state* to *goal-gradient* (used in the learning phase of an oscillatory cycle) was removed and the *goal-gradient* received steady input from the *goal* layer, interrupted only by the state-change inhibition signal. In other words, the *goal-gradient* layer continuously represented the actual goal gradient rather than shifting into learning mode half of the time.

In the task, participants first saw an information screen from which they could determine the immediate goal state and the appropriate next action. This plan was maintained over a delay before being implemented. At the beginning of each trial simulation, the queuing module was set to "load" while the model interactions determined the best method of getting from the current state to the goal state. This period is analogous to the period in which subjects look at the starting information screen and plan their next move. Then, the queuing module was set to "execute," modeling the period in which participants are prompted to make their selection. Finally, the chosen action implements a state transition and the environment provides new state information to the *state* layer, modeling the outcome phase of the experiment.

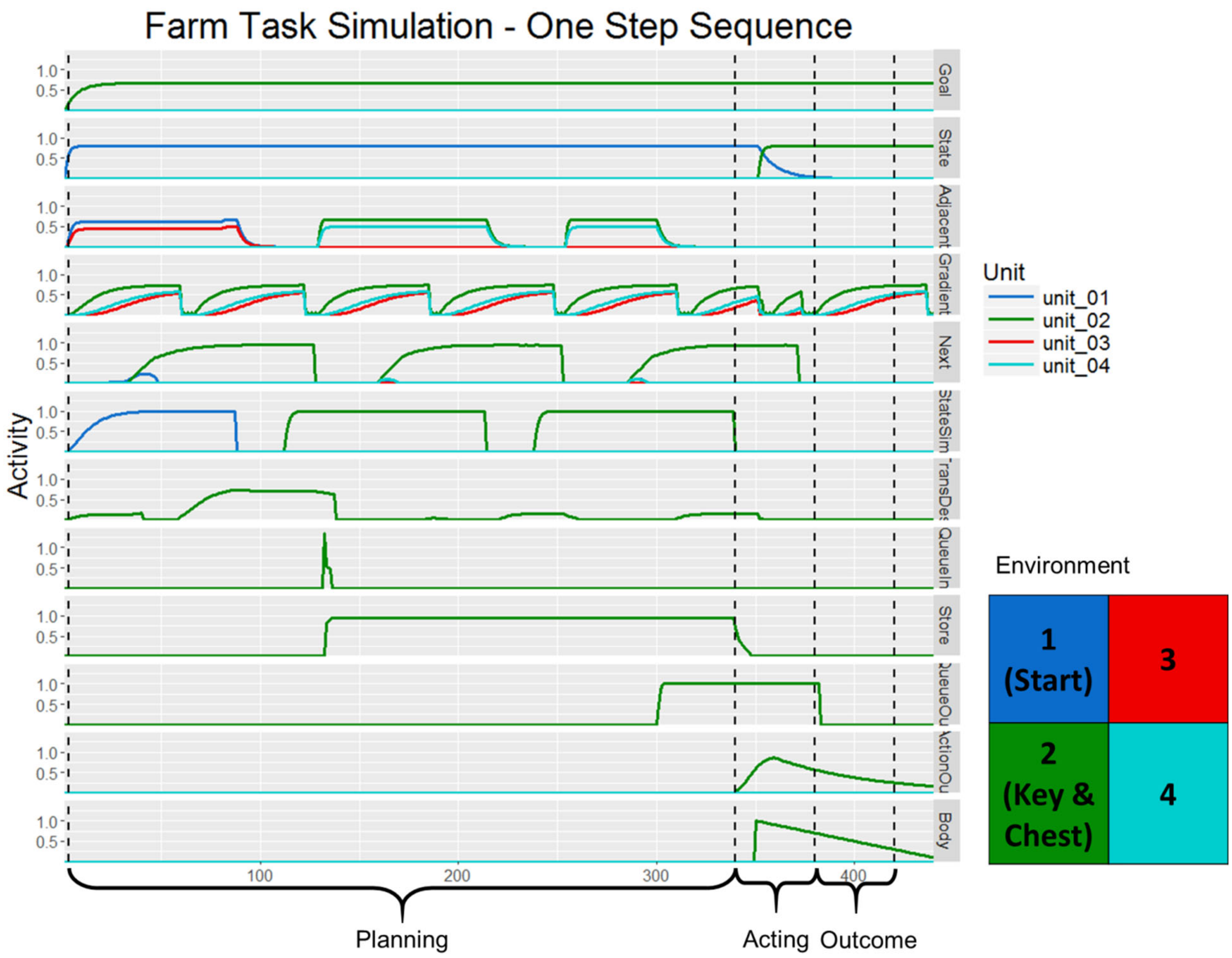


**Figure S31. Model activity during a simulated one-step sequence of the Treasure Hunt task.** The competitive queueing module first loads a plan and then executes it sequentially. State activity shows that the agent remains in state 1 for the first half of the simulation, while simulated-state (StateSim) shows the state transition the agent simulates as it forms its plan. Adjacent-states (Adjacent) receives input from stateSim which, along with goal-gradient (Gradient) activity determines the desired next state and therefore the appropriate transition to make. The plan is kept in queue-store (Store) which receives a burst of input from queue-input (QueueIn) and finally executes the plan by sending output to queue-output (QueueOut) which drives the motor system. The vertical dashed lines indicating the different phases of the simulation used in the creation of the model RDMs. For each layer, activity within each period was averaged across time to form a single vector representing the average pattern for that time period in the trial type being simulated. The bounds of each phase were determined qualitatively. The planning period is longer than the acting and outcome periods because the model takes longer to form a plan than execute it or observe the outcome.

Some pairs of trials in the task comprised a two-step sequence in which the final goal was initially two states away from the starting state. On the second trial of such sequences, participants were not provided any information on the information screen at the start of the trial, ensuring that they had encoded and maintained all goal-related information from the information screen presented at the start at the first trial in the sequence. These pairs of trials were modeled within a single GOLSA simulation. The model seeks the quickest path to the goal, identifying immediately available subgoals as needed. However, in the task, the location of the key necessitated a specific path to reach the final goal of the treasure chest. To provide these instructions to the model at the start of a two-step simulation, the goal representation from the subgoal (the key) was provided to the model first until the appropriate action was loaded and then the goal representation shifted to the final goal (the chest). Once the full two-step state transition sequence was loaded in the queue, the actions were read out sequentially, as shown in Figure S4.

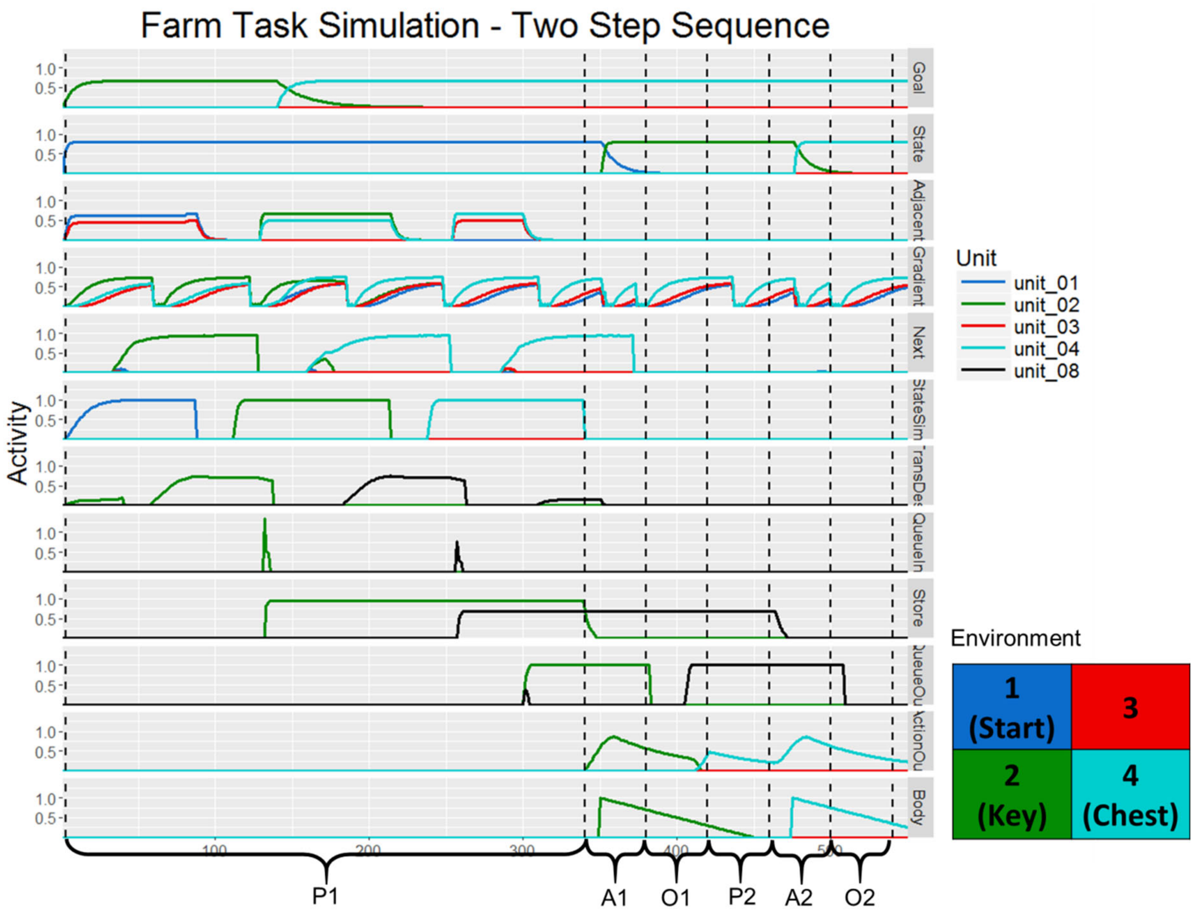


**Figure S4. Model activity during a simulated two-step sequence of the Treasure Hunt task.** The competitive queueing module first loads a plan and then executes it sequentially. State activity shows that the agent remains in state 1 for the first half of the simulation, while simulated-state shows the state transitions the agent simulates as it forms its plan. Adjacent-states receives input from simulated-state which, along with goal-gradient activity determines the desired next state and therefore the appropriate transitions to make. The plan is kept in queue-store which receives bursts of input from queue-input and finally executes the plan by sequentially sending output to queue-output which drives the motor system. To force the agent to go to the appropriate intermediate state, goal activity first reflects the key location and then the chest location. The vertical dashed lines indicate time periods used when creating the RDMs for the two-step sequence simulations. The first three time periods correspond to the first trial in the sequence while the latter three correspond to the second trial in the sequence. Again, the first planning period is much longer due to the nature of the model dynamics. During the second "planning" period (P2), the plan was already formed as must have been the case in the actual experiment since on the second trials in a two-step sequence, no information was presented at the start of the trial and had to be remembered from the previous trial.

A separate RDM was generated for each model layer. Patterns were extracted from three time intervals per action (six total for the two-step sequence simulations). Due of the time required to load the queue, the first planning period was longer than all other intervals. For each simulation and time point, the patterns of activity across the units were averaged over time, yielding one vector. Each trial type was repeated 10 times and the patterns generated in the previous step were averaged across simulation repetitions. The activity of each layer was thus summarized with at most 96 patterns of activity which were converted into an RDM by taking one minus the Pearson correlation between each pattern. Patterns in which all units were 0 were ignored since the correlation is undefined for constant vectors.

We looked for neural regions corresponding to the layers that played a critical role in the model during the acting phase in the typical learning oscillation since in these simulations the learning phase of the oscillation was disabled. We created RDMs from the following layers: *current-state*, *adjacent-states, goal*, *goal-gradient, next-desired-state*, *desired-transition*, *action-out*, *simulated-state*, and *queue-store*. As a control, we also added a layer component which generated normally distributed noise ($\mu = 1$, $\sigma = 1$).

**Searchlight**

The searchlight analysis was conducted using Representational Similarity Analysis Toolbox, developed at the University of Cambridge (http://www.mrc-cbu.cam.ac.uk/methods-and-resources/toolboxes/license/). For each of these layer RDM, a searchlight of radius of 10mm was moved through the entire brain. At each voxel, an RDM was created by from the patterns in the spherical region centered on that voxel.

An r value was obtained for each voxel by computing the Spearman correlation between the searchlight RDM and the model layer RDM, ignoring trial time periods in which all model units showed no activity. A full pass of the searchlight over the brain produced a whole-brain r map for each subject for each layer. Voxels in regions that perform a similar function to the model component will produce similar RDMs to the model component RDM and thus will be assigned relatively high values. The r maps were then Fisher-transformed into z maps ($z = \frac{1}{2}\ln\left(\frac{1+r}{1-r}\right)$). The z maps were normalized into the MNI template and smoothed with a Gaussian kernel (FWHM = 8 mm). Second level effects were assessed with a t test on the normalized and smoothed z maps, with a cluster defining threshold of $p < 0.001$, cluster corrected to $p < 0.05$ overall. The complete results are shown in Table S3.

### RSA results

| | | MNI Coordinates | | | | | |
|---|---|---|---|---|---|---|---|
| **Layer** | **Peak Region (TD label)** | **X** | **Y** | **Z** | **Z-Score** | **p** | **Size** |
| goal | Cuneus | -14 | -86 | 17 | 6.75 | <0.001 | 16019 |
| goal-gradient | Cuneus | 14 | -83 | -3 | 7.63 | <0.001 | 13618 |
| goal-gradient | Postcentral Gyrus | -21 | -38 | 68 | 4.25 | 0.01 | 192 |
| goal-gradient | Superior Frontal Gyrus | -28 | 45 | 37 | 3.7 | 0.025 | 149 |
| adjacent-state | Middle Temporal Gyrus | 65 | 0 | -31 | 3.82 | 0.014 | 164 |
| next-desired-state | Middle Frontal Gyrus | 17 | 34 | -14 | 5.09 | <0.001 | 3576 |
| next-desired-state | Lentiform Nucleus | 28 | -10 | 14 | 3.96 | 0.025 | 161 |
| desired-transition | Sub-Gyral | -24 | -10 | 37 | 6.41 | <0.001 | 7494 |
| action-output | Precentral Gyrus | -38 | 0 | 37 | 5.24 | 0.001 | 379 |
| action-output | Middle Frontal Gyrus | 31 | -7 | 41 | 4.98 | <0.001 | 1030 |
| action-output | Supramarginal Gyrus | -38 | -45 | 37 | 4.21 | 0.048 | 142 |
| queue-store | Superior Frontal Gyrus | 17 | 65 | 27 | 4.2 | 0.02 | 224 |
| queue-store | Middle Temporal Gyrus | 62 | 3 | -31 | 4.16 | 0.002 | 436 |
| simulated-state | Lingual Gyrus | 3 | -79 | -7 | 5.16 | <0.001 | 1768 |
| simulated-state | Middle Temporal Gyrus | 45 | 28 | -34 | 4.38 | <0.001 | 1225 |
| simulated-state | Middle Frontal Gyrus | -52 | 45 | -3 | 3.81 | 0.001 | 388 |
| state | Extra-Nuclear | -31 | -14 | 24 | 7.84 | <0.001 | 46188 |

**Table S2.** Significant similarity clusters for RSA analysis. The p and Size columns refer to cluster-corrected values. Anatomical labels are derived from the Automated Anatomical Labeling Atlas in SPM5 (Tzourio-Mazoyer et al., 2002).
